# Fish metabolome from sub-urban lakes of the Paris area (France) and potential influence of noxious metabolites produced by cyanobacteria

**DOI:** 10.1101/2020.12.14.422619

**Authors:** Benjamin Marie, Alison Gallet

## Abstract

The recent democratization of high-throughput molecular phenotyping allows the rapid expansion of promising untargeted multi-dimensional approaches (*e.g.* epigenomics, transcriptomics, proteomics, metabolomics, …). Indeed, these emerging omics tools, processed for ecologically relevant species, may present innovative perspectives for environmental assessments, that could provide early warning of eco(toxico)logical impairs. In a previous pilot study (Sotton et al., Chemosphere 2019), we explore by ^1^H NMR the bio-indicative potential of metabolomics analyses on the liver of 2 sentinel fish species (*Perca fluviatilis* and *Lepomis gibbosus*) collected in 8 water bodies of the peri-urban Paris’ area (France). In the present study, we further investigate on the same samples the great potential of high-throughput UHPLC-HRMS/MS analyses. We show that the LC-MS metabolome remarkably allows clear separation of individuals according to the species, but also according to their respective sampling lakes. Interestingly, similar variations of *Perca* and *Lepomis* metabolomes occur locally indicating that site-specific environmental constraints drive the metabolome variations beyond the obvious genetic differences between the two species, and seem to be influenced by the production of noxious metabolites by cyanobacterial blooms in certain lakes. Thus, the development of such reliable environmental metabolomics approaches is constituting an innovative bio-indicative tool for ecological stress assessment, such as toxigenic cyanobacterial blooms, and aim at being further follow up.

## 1. INTRODUCTION

In the past decades, the use of bio-indicator aquatic organisms has been widely adopted to monitor and to manage the quality of water bodies worldwide (Lorenz 2003), together with a growing recognition of the difficulties and the rational limits of direct chemical monitoring for providing sufficient information to adequately assess the risks from anthropogenic chemicals in the natural environment (Reyjol et al., 2014). More recently, the recent democratisation of high-throughput molecular phenotyping has allowed the rapid expansion of promising untargeted multi-dimensional approaches (*e.g.* epigenomics, transcriptomics, proteomics, metabolomics), constituting as much as responding biological traits, that now represent promising perspectives for environmental assessments (Bahamonde et al. 2016; Cordier et al., 2020). It is very likely that emerging omics-based analyses, such as metabolomics, developed for ecologically relevant aquatic species, may support the development of new ecological evaluation tools, that could provide even earlier warning of eco(toxico)logical impairs (Marie 2020).

The metabolome, defined as the set of primary metabolites synthesized at a given time and thus represents its global metabolic chemical picture, is assumed to represent a final endpoint of the phenotypic response of an organism that can potentially be modified when the ecological and environmental stress conditions change (Bundy et al., 2008). In this way, metabolomic studies have become a relevant approach to describe and analyse physiological processes involved in the homeostatic responses of the organisms encountering environmental stresses from potentially multiple origins (Capello et al., 2016; Good et al., 2020). However, despite its high potential to better understand the molecular mechanisms implicated in the ecotoxicological responses of aquatic organisms (Sardans et al., 2011; Viant, 2008), such environmental metabolomics investigations remains still rare.

Fishes, in particular, present unique features that make them especially relevant for bio-indication purposes, as they occupy almost any aquatic habitat, being capable of experiencing different and variable environmental conditions (Harris 1995). Fishes are also responsive to numerous abiotic (temperatures, water velocities, sediment loads, hypoxia, …) or biotic (famine, predation, parasitism, …) pressures, together with anthropogenic stressors, such as contaminants, that represent additional constraints that fish may experience in disturbed ecosystems (McClanahan 2019). Environmental stressors are known to induce biological variations, concerning the specific physiological, developmental and reproductive capabilities of the different fish species allowing them to colonize and occupy ecosystems (Hamilton et al., 2016).

As cyanobacterial blooms are one of the most common stress encountered by organisms living in modern freshwater lentic systems around the world (Harke et al., 2016; Burford et al., 2020), several recent efforts have been attempted to characterize the potential effect of cyanobacteria proliferation on various fish species according to experimentation performed in the lab, in mesocosms, or from field sampling (Amado et al., 2010; Ferrao-Filho et al., 2011; Sotton et al., 2017a, 2017b; Le Manach et al., 2018). To this end, we initially investigated the liver metabolome of fish collected from different lakes encountering low or high cyanobacterial proliferation by ^1^H NMR technics (Sotton et al., 2019). This pilot study leads to the description of putative metabolite concentrations, corresponding to specific NMR signals, that were significantly correlated with higher cyanobacterial occurrence.

The present study investigates the bio-indicative potential of high-throughput molecular phenotyping supported by LC-HRMS approaches on 2 sentinel fish specimens (*Perca fluviatilis* and *Lepomis gibbosus*) collected from a set of 8 water bodies of the peri-urban area of Paris (France), that were previously investigated by ^1^H NMR (Sotton et al., 2019). We hypothesis that LC-MS-based metabolomics approach may present a higher potential for environmental diagnosis, as providing valuable multi-metric characters indicative of biological or ecological conditions, together with a prospective mechanistic under-standing that supports cause-effect relationship investigation. With regard to the cyanobacterial occurrence and the associated production of bioactive metabolites in the different lakes, we compare here the metabolome of two different sentinel species. Indeed, these organisms could present different or comparable sensitivity and responsiveness to local eco(toxico)logical constraints (*i.e.* potentially the production of noxious metabolites by cyanobacteria), supporting the general interest for omics-based environmental approach of cyanobacterial bloom threat.

## 2. MATERIALS AND METHODS

### 2.1. Fish sampling

Field sampling campaigns was performed during late summer 2015 (7-10^th^ September) in 8 peri-urban pounds around Paris’ area (Île-de-France region, France), chosen for their respective eutrophication levels and the presence or the absence of recurrent cyanobacterial blooms: Cergy-Pontoise (Cer), Champs-sur-Marne (Cha), Maurepas (Mau), Rueil (Rue), Verneuil (Ver), Varennes-sur-Seine (Var), Fontenay-sur-Loing (Fon) and Triel (Tri) pounds (Supp. figure S1) (Maloufi et al., 2016; Sotton et al., 2019). These sites were sampled by the Hydrosphère company (www.hydrosphere.fr) with electric fishing device (FEG 8000, EFKO, Leutkirch, Germany) for capturing fish alive, that were then promptly sacrificed for liver collection. The investigation of the fish guild indicates that only the perch (*Perca fluviatilis*) and pumpkinseed sunfish (*Lepomis gibbosus*) were presents in all or almost all these pounds (supplementary table S1) and were further selected as sentinel species for further molecular phenotyping analyses by metabolomics.

Briefly, alive caught fishes (n=5-10 young-of-the-year per pounds and per species) were directly measured (12.0±4.8 cm), weighed (9.3±2.6 g), briefly euthanized by neck dislocation and then liver of each individual was shortly sampled, flash-frozen in liquid nitrogen and kept at −80°C until analyses, in accordance with European animal ethical concerns and regulations. In every lake, sub-surface chlorophyll-*a* equivalent concentrations attributed to the four-main phytoplankton groups (Chlorophyta, Diatoms, Cyanobacteria and Cryptophyta) were measured with an *in-situ* fluorometer (Fluoroprobe II, Bbe-Moldenke, Germany) (Supplementary figure S2). Sub-surface water samples filtered on 20-*μ*m mesh size were also collected for phytoplanktonic community analysis and further metabolomics characterisation, and then kept at −80°C until analysis as previously described (Sotton et al., 2019).

### 2.2. Liver metabolite extraction and metabolomics analyses

The liver extraction was performed on 132 individuals (comprising 78 *Perca* and 54 *Lepomis*) with methanol/chloroform/water (ratio 2/2/1.8 – 22 mL.g^−1^ at 4°C) and the polar fraction was analyzed on a 600-MHz NMR spectrometer equipped with a 5-mm cryoprobe (Advance III HD Bruker, Germany) with a noesygppr1d pulse sequence as previously described (Sotton et al., 2019). ^1^H-NMR spectra were treated with Batman R-package for deconvolution, peak assignment and quantification of 222 putative metabolites (Hoa et al., 2014).

The liver extracted polar phase was additionally injected (2 μL) on C_18_ column (Polar Advances II 2.5 pore - Thermo), then eluted at a 300 μL.min^−1^ flow rate with a linear gradient of acetonitrile in 0.1% formic acid (5 to 90 % in 21 min) with an ultra-high-performance liquid chromatography (UHPLC) system (ELUTE, Bruker). Consecutively, the individual metabolite contents were analysed using an electrospray ionization hybrid quadrupole time-of-flight (ESI-Qq-TOF) high-resolution mass spectrometer (Compact, Bruker) at 2 Hz speed on positive MS mode on the 50–1500 *m/z* range. The feature peak list was generated from recalibrated MS spectra (< 0.5 ppm for each sample, as an internal calibrant of Na formate was injected at the beginning of each sample analysis) within a 1-15 min window of the LC gradient, with a filtering of 5,000 count of minimal intensity, a minimal occurrence in at least 50% of all samples, and combining all charge states and related isotopic forms using MetaboScape 4.0 software (Bruker).

Additionally, five pools of six different individuals randomly selected for *Perca* and *Lepomis* (quality check samples) and phytoplanktonic biomass extracts from the 8 lakes were similarly eluted then analysed on positive autoMS/MS mode at 2-4 Hz on the 50-1500 *m/z* range for further metabolite annotation. Molecular networks were performed with GNPS (http://gnps.ucsd.edu) and/or MetGem (http://metgem.github.io) softwares, as previously described (Kim Tiam et al., 2019; Le Manach et al., 2019) for cyanobacterial and fish metabolite annotation was assayed using in-house (Le Manach et al., 2019) or CyanoMetDB (Jones et al., 2021), and GNPS, Mona, HMDB and Massbank MS/MS spectral libraries. Water samples of each lake, concentrated on a 20-*μ*m mesh size, were also extracted with 75% methanol (2 min sonication, 5 min centrifugation at 15,000 g - 4°C) and then similarly analysed in triplicates on LC-HRMS on autoMS/MS positive mode, as earlier described, and metabolite list annotated with same pipelines.

### 2.4. Data matrix treatment

The resulting the intensity data tables of the 222 metabolites (^1^H NMR) and 1252 analytes (LC-MS) were further treated for quantile normalization and Pareto’s scaling, inspected data representation by PCA and heatmap with hierarchical clustering (Euclidean distance) and then analysed to investigate the influences of “Species” and “Lakes” parameters on the datasets by PERMANOVA or PLS-DA using MixOmics R Package (Rohart et al., 2017), MetaboAnalyst 5 (Chong et al., 2019) and MicrobiomeAnalyst tools (Chong et al., 2020).

## 3. RESULTS AND DISCUSSION

### 3.1. Distinction of *Perca* and *Lepomis* liver LC-MS metabolome

Figure 1 illustrates the global relative metabolites analyzed by LC-MS for each fish (comprising *Perca* and *Lepomis*) on a heatmap with hierarchical classifications and individual plot principal component analyses (PCA). A very strong and significant discrimination according to the “species” was achieved for this LC-MS liver metabolomics dataset (PERMANOVA F-value = 94.92; R^2^ = 0.43; P-value <0.001), when a slighter but still important discrimination can be also retrieved for “lakes” (PERMANOVA F-value = 4.96; R^2^ = 0.22; P-value <0.001).

**Figure 1.**
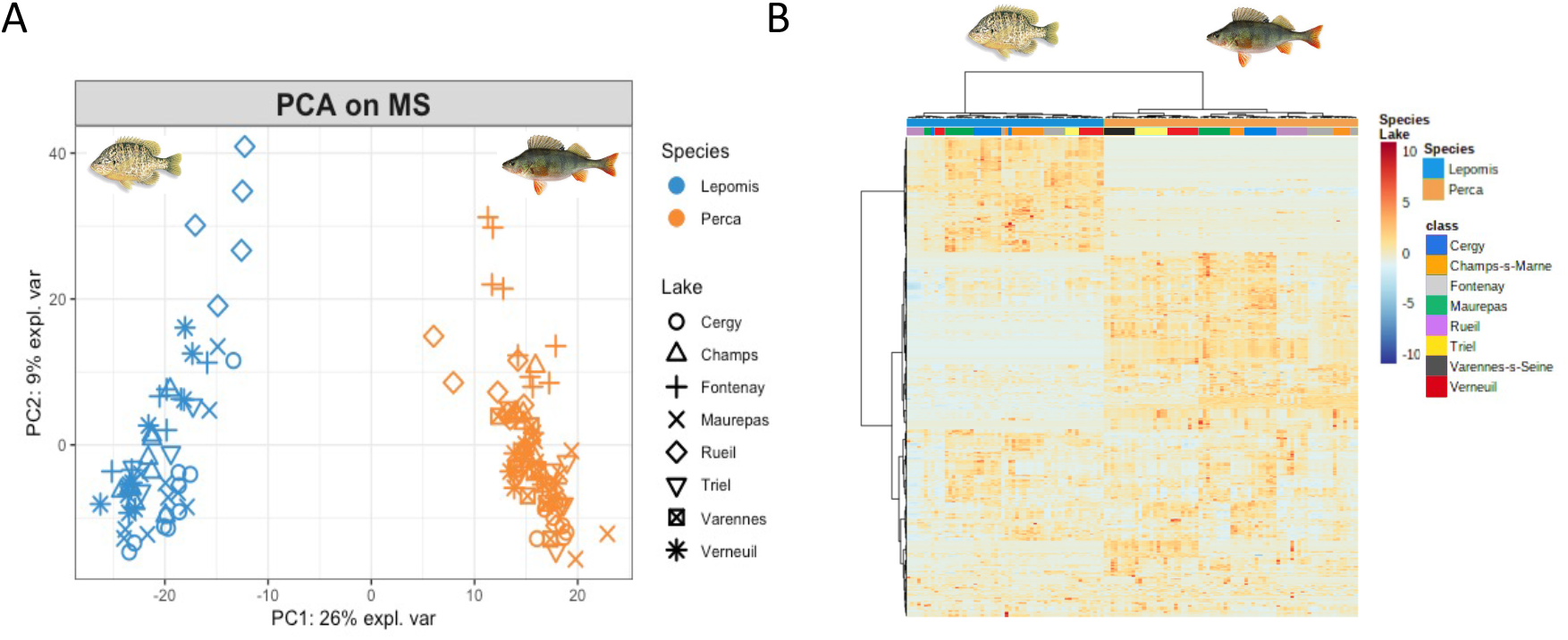
Visualization of the dataset structuration for LC-MS metabolomics (comprising 132 fish and 1252 analytes) on PCA (A) and heatmap with hierarchical clustering (B) for *Perca* (n=78) and *Lepomis* (n-54) fish collected during the 7-10^th^ of September 2015 within 8 pounds of the peri-urban Paris’ area.

In parallel, a molecular network performed with fragmentation data was obtained by LC-MS/MS analyses of *Perca* and *Lepomis* metabolomes. This approach supports the global annotation process of the fish liver metabolites considering both structural identity and similarity (Nothias et al., 2020). The GNPS and the *t*-SNE representation of the molecular network highlights that most, if not all, of the known metabolites (presenting structural identity or analogy hits) are largely shared between the two species (*e.g.* nucleic acids, carnitines, glutathiones, lipids, saccharides, …). Moreover, most of the species-specific cluster metabolites remain uncharacterized, as corresponding molecules and clusters present no match within the public chemical databases (supplementary figure S2) (da Silva et al., 2015). Taken together, these analyses remarkably illustrate the various specificities of the metabolome of these two species. It also shows the substantial portion of the specific liver molecular metabolism still uncharacterized and that could be related to species-specific nutritional, physiological or toxicological capabilities (Brusle et al., 2017).

### 3.2. Comparison of *Perca* and *Lepomis* LC-MS metabolome between lakes

In addition, the LC-MS dataset of the two fish species considered separately clearly show even more consistent discrimination of the metabolomes of fish originating from the different sampling lakes. The un-supervised PCA and the heatmap with hierarchical classification (Figure 2), performed with LC-MS dataset of liver metabolomes, present distinct local signatures that support reliable discrimination of the lake of sampling for both *Perca* and *Leptomis* (Fig. 2A–B and 2C–D, respectively). These observations strongly suggest that the locality seems to globally influence the LC-MS metabolome composition for both species. However, the liver metabolomes of individual *from* Fontenay-sur-Loing lake (Fon) exhibite a quite species-specific relative positions on PCA, considering the relative position of individual metabolomes from other lake in the two species (Figure 2A and 2C). This suggests that, although for most localities the specific “lake” signature of the metabolome appears in good agreement between the two species, in some specific environments, the metabolome signatures are variable from one species to the other one. The species-specific signature of the *Perca* and *Lepomis* metabolomes in Fon lake potentially traduce rather specific responsiveness and/or sensitivity to a local factor.

**Figure 2.**
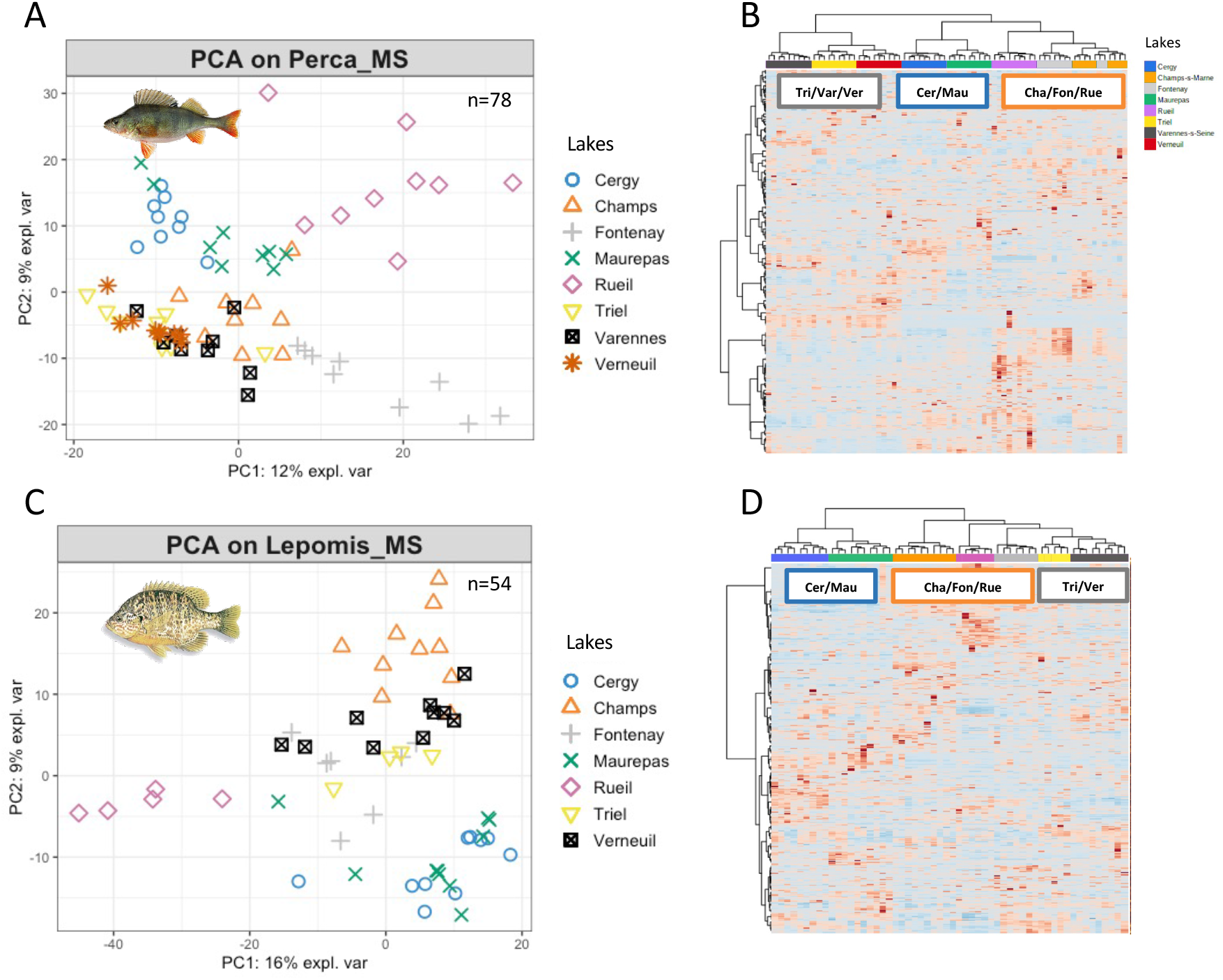
Individual plot of PCA (A and C) and heatmap with hierarchical classification (B and D) of LC-MS liver metabolomes of *Perca* (n=78; A-B) and *Lepomis* (n=54; C-D) collected during the 7-10^th^ of September 2015 within 8 pounds of the peri-urban Paris’ area.

Interestingly, the hierarchical classifications performed respectively on LC-MS metabolome datasets of *Perca* and *Lepomis* show very similar lake relationships, grouping together the fish from the lakes of Cer and Mau, from Cha, Fon and Rue, and from Tri, Var and Ver for *Perca* or Tri and Ver for *Lepomis*, as this latter was not retrieved in Var (Figure 2B and 2D). This observation suggests that these environments could present comparable, if not homogenous, environmental conditions. Then, this hypothesis was further considered and potential explicative factors investigated, considering the potential influence of cyanobacterial proliferation.

Indeed, as previously shown, the Fon, Tri, Var and Ver lake encounter a remarkable high amount of cyanobacteria belonging to the *Planktothix*, *Pseudo-Anabaena*, *Aphanizomenon*, *Anabaena* or *Microcystis* genera, when others (Cer, Mau, Cha and Rue) exhibit mostly green algae, diatoms or cryptophytes (Fig. 3A–B). Interestingly, a molecular network performed with LC-MS/MS data obtained from the respective phytoplanktonic biomass of these lakes shows that noticeable amounts of cyanopeptides are observed in the water column of the cyanobacterial-rich lakes (Fon, Tri, Var and Ver), with the specific presence of most noxious cyanopeptides (microcystins, cyanopeptolins and anbaenopeptins) being detected in Tri, Var and Ver lakes, when Fon exhibits rather less toxic cyanopeptides (Janssen 2019), such as microginins and aeruginosins (Fig. 3C). Taken together, these observations suggest that the proliferation and the subsequent production of these potentially noxious cyanobacterial metabolites could be considered as an explicative factor for the local metabolome singularity of the liver metabolome of *Perca* and *Lepomis* from Tri, Var and Ver lakes, as previously proposed (Sotton et al., 2019).

**Figure 3.**
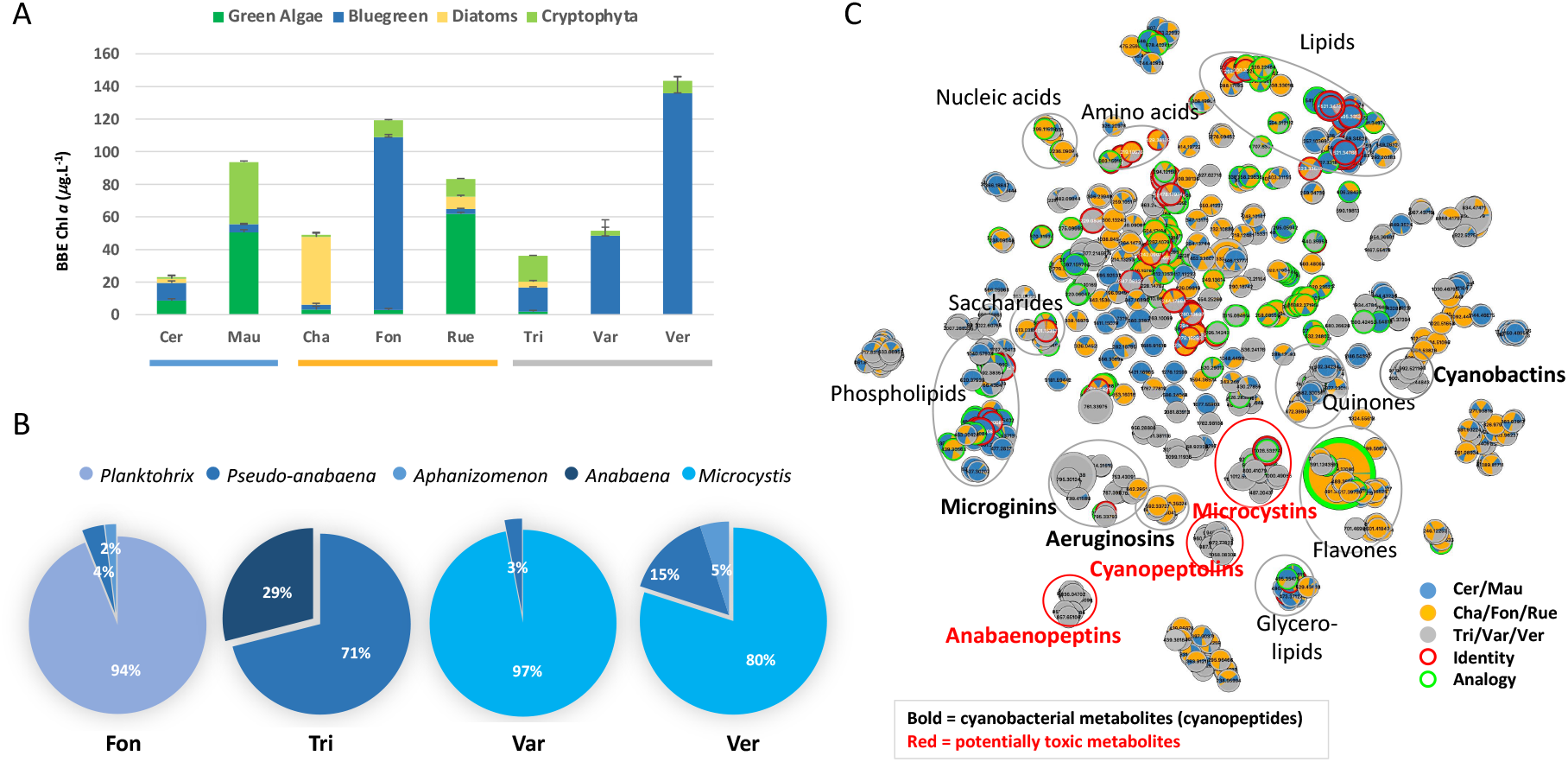
Phytoplankton composition estimated by BBE measurment of the 8 lake sub-surface water (A), corresponding cyanobacteria relative composition for Fon, Tri, Var and Ver (B), and molecular networking of metabolites extracted from the filtered biomass of the respective water of the 8 lakes generated with *t*-SNE algorithm, with cyanobacteria peptide clusters indicated in bold and potentially more noxious family clusters (microcystin, cyanopeptolmin and anbaenopeptin) indicated in red (C).

### 3.3. Most discriminating metabolites between the different lake groups

Thus, we re-explore the fish metabolome discrimination considering together fish from lake presenting noxious cyanobacterial metabolites (“Tri/Var/Ver” group for *Perca* and “Tri/Ver” group for *Lepomis*) or not (“Cer/Mau/Cha/Fon/Rue” group), by PLS-DA to identify metabolite that the most discriminate between these two lake groups (Figure 4). Indeed, while unsupervised PCA were first used to evaluate the global dispersion between species and sampling groups, a supervised model such as partial least square differential analysis (PLS-DA) allows to maximize the separation between sample classes (*e.g.* “lakes”) and to extract information on discriminating features (variable importance on projection - VIP). For both fish species the metabolomes present a consistent discriminant analysis scoring (*Perca*: accuracy = 1, performance R^2^ = 0.78, predictability Q^2^ = 0.72; *Lepomis*: accuracy = 0.93, R^2^ = 0.59, Q^2^ = 0.29) and large set of discriminating variables (*Perca* VIP_>1.5_=109 and *Lepomis* VIP_>1.5_=93) (Figure 4A–B). Among these metabolites, 33 components have been successfully annotated considering the molecular mass, the isotopic pattern, together with the fragmentation pattern of all components with GNPS and MetGem algorithms and manual curation. Overall, the observation of the metabolite semi-quantification of the annotated best VIP presents largely similar variations between these two groups of lakes for *Perca* and *Lepomis* (Figure 4C; Supp. fig. S3). Indeed, except for the reduced-glutathione that present higher concentrations in the livers of *Perca* from “Tri/Var/Ver” lakes, when livers of *Lepomis* exhibits lower concentrations than in the those of fish of the “Cer/Mau/Cha/Fon/Rue” group, all other metabolites present similar pattern in the two species. This discrepancy suggests that glutathione-dependent redox mechanisms could be more extensive in *Perca* than in *Lepomis* livers.

**Figure 4.**
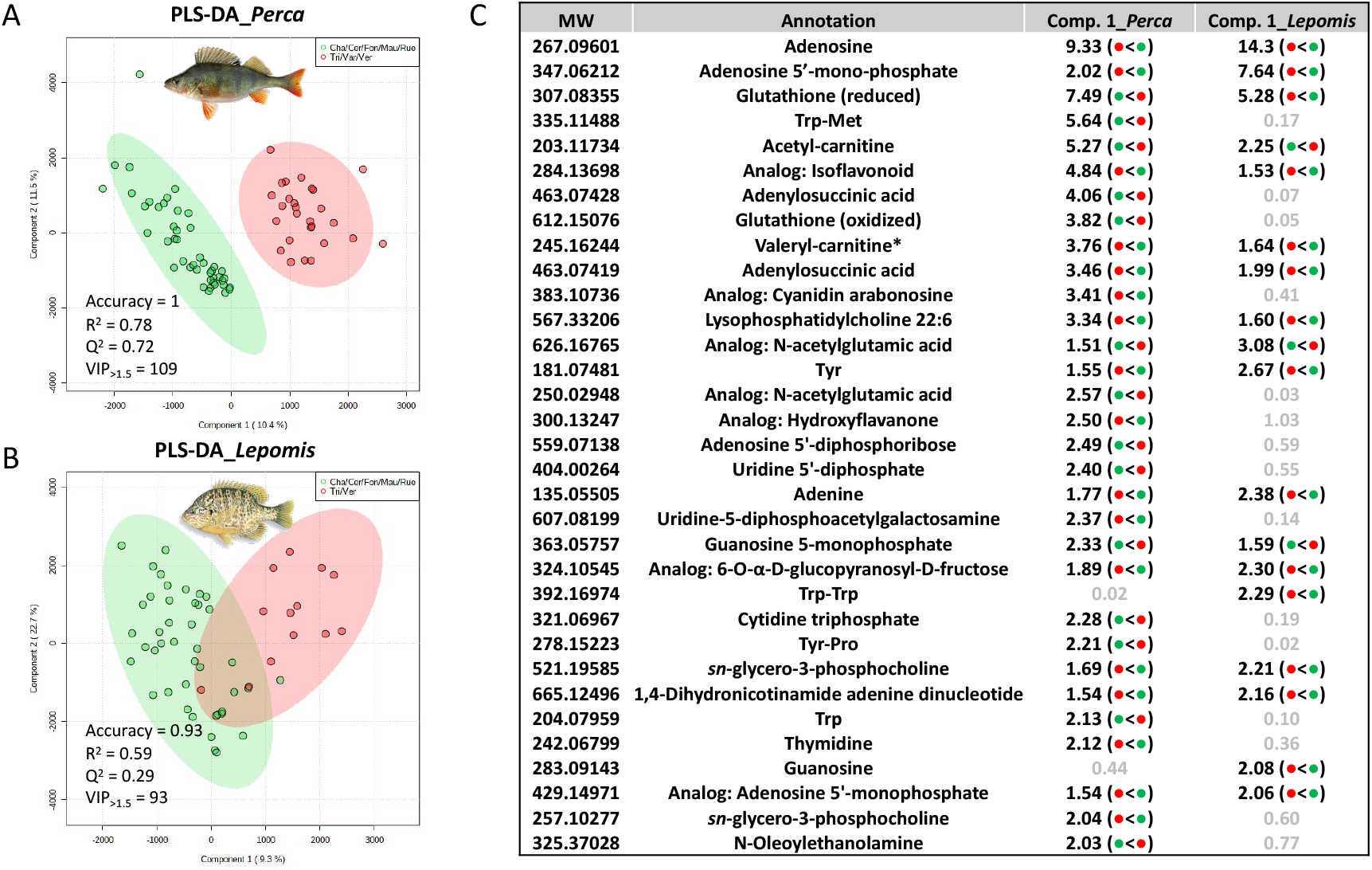
Individual plots of PLS-DA, testing the difference between the different lake groups presenting, or not, high content of potentially noxious cyanobactérial metabolites (Cha/Cer/Fon/Mau/Rue and Tri/Var/Ver, respectively) according to the LC-MS metabolomes of *Perca* (A) and *Lepomis* (B), and corresponding list of annotated best VIP list (metabolite score > 1.5) of *Perca* and *Lepomis* metabolomes (C). Accuraty, predictability and quality performances and number of VIP metabolite score > 1.5) for sampling lake discrimination are indicated for each species. Dots between brackets indicate when the metabolite concentration is more important in the Cha/Cer/Fon/Mau/Rue (green) or Tri/Var/Ver (red) lake group. * indicates that this metabolite was also observed as being negatively correlated with cyanobacteria occurrence within the different lake according to NMR analysis (Sotton et al., 2019).

Globally, the main differences between the two lake groups of the different metabolite concentration show that fishes collected from the “Cer/Mau/Cha/Fon/Rue” seem to present greater energetic, anti-oxidant/detoxification and lipids reserves (*e.g.* various nucleic acids, glutathiones, flavonoids and lipo-phosphocholines) than in lakes presenting more cyanobacteria and related noxious metabolites. In addition, greater amounts of amino acids, di-peptides and acylcarnitines could be related to higher protein catabolism and beta-oxidation in the liver of fish originating from the “Tri/Var/Ver” lakes, hypothetically traducing higher energetic requirements for those organisms. Taken together, these observations are in agreement with the hypothesis of the occurrence of higher stress conditions locally occurring in the Tri, Var and Ver lakes, in correspondence with the local production of potentially noxious metabolites by cyanobacterial proliferation (Sotton et al., 2017a, 2017b; Le Manach et al., 2018).

In general, various genetic or phenotypic factors (such as development stages, contamination levels, predator/parasite pressures, food availability, …) could influence and explain lthe ocal discrepancy of the metabolome of fishes collected from different environments (reviewed in Marie 2020). However, in the present case, the fact that *Perca* and *Lepomis* metabolomes analysed in parallel present similar metabolite variations clearly indicates that local environmental constraints drive such phenotypic co-variations (beyond the obvious genetic distinction between *Perca* and *Lepomis*). Then, these present results claim the great value of LC-MS-based metabolomic imprints for molecular phenotyping on sentinel species and the exploration of environmental stresses (Lohr et al., 2019).

### 3.4. Comparison of ^1^H NMR and LC-MS metabolomics for fish livers

Considering the previous ^1^H NMR metabolomics datasets of these two fish species retrieved from the same samples (Sotton et al., 2019), the NMR liver metabolome of *Perca* and *Lepomis* shows more limited specificity according to “species” or “lakes” factor than LC-MS dataset. Although the 222 potential metabolites quantified by NMR on 132 fish (78 *Perca* and 54 *Lepomis*) present no obvious structuration of the individual dataset according to “species” or “lakes” variables when observed on PCA or hierarchical classification, slight but significant relations are however supported by PERMANOVA (F-values = 4.15 and 3.63; R^2^ = 0.03 and 0.17; P-values < 0.01 and 0.01, for “species” and “lakes” respectively, Supp. fig. S3). Indeed, according to the relatively lower performances of the hierarchical classifications and PERMANOVA and the PLS-DA (Supp. fig. S4 and S5), the NMR liver metabolomes present restricted discrimination on “lakes” for both species and could support here only a limited discrimination potential, when compared to LC-MS metabolomics performances.

Overall, the dataset obtained from the LC-MS metabolome analysis presents a clearer discriminant pattern between the two species or the 8 lakes, than the NMR dataset generated on the same samples (with the same extraction procedure), regarding both hierarchical classification, individual plots of PCA or PERMANOVA performances (Fig. 1 and 2; Supp. fig. S4 and S5). This observation is further confirmed by direct comparison of the NMR and LC-MS metabolome dataset using MixOmics Diablo tool for multi-block PLS-DA (Supp. fig. S6 and S7). Indeed, although the two datasets appear mostly to be congruent, the individual plot distribution carries much more variance on the PLS-DA projection for the LC-MS analysis than for the NMR, considering both fish species.

On one side, the ^1^H NMR metabolomics allows the reliable quantification of main liver metabolites (belonging mostly to amino acids, sugars, TCA metabolites, …), by detecting characteristic chemical groups, with regards to their respective position within the molecules (Martin et al., 2015). These primary metabolites are key contributors to the cellular metabolism and are supposed to be responsive to variable physiological conditions of fish encountering peculiar biological outcomes and/or environmental constraints (Roques et al., 2020). However, this approach remains focused on few principal metabolites involved in main cellular pathways which measured cellular contents (in *ppm*) can be involved in multiple regulation processes, and not being specific of subtle molecular regulation processes. On the other hand, ^1^H NMR presents only low sensitivity and limited discrimination capability for molecules presenting similar chemical structures, such as lipids, that exhibits an aggregated and unspecific signal on NMR spectra (Emwas et al., 2019). For this reason, it is supposed to present lower discriminating power than more selective and less qualitative, but more sensitive, high-resolution mass spectrometry-based metabolomics approaches.

On the other side, although LC-MS-based metabolomic remains a selective analytical approach (according to both LC separation and ionisation respective properties of each molecule), it presents un-precedent analytical capabilities in terms of sensitivity, dynamic range and number of characterized components and offer now unprecedented perspectives for the investigation for molecular phenotyping applied to environmental sciences (Beyoglu and Idle., 2020). We assume, that a single LC-MS fingerprint analyse does not embed the whole metabolite picture of the biological compartment (the fish liver, in the present case), especially because of its selectivity performances (Gika et al., 2019). However, it provided specific and precise measurements of a large number of components that can endorse high-throughput and in-depth phenotyping. For comparison, ^1^H NMR metabolomics allows the global quantification of the CH_2_-CH_2_ or CH_2_-CH=CH fatty acid bounds (Marchand et al., 2018), when LC-MS metabolomics can discriminate and characterize hundreds of different lipids belonging to various sub-classes (Dreier et al., 2020), providing undoubtedly more informative and discriminant features. In addition, multi-omics integration tools, such as MixOmic DIABLO (Singh et al., 2019), provides innovative solutions for the comparison of heterogeneous dataset matrices obtained from a single set of samples or individuals (supplementary figures S6-S7).

### 3.5. Use of LC-MS metabolomics fingerprint for environmental assessment?

Although the value of LC-MS metabolomics for investigating the impacts of environmental stressors or contaminants, and their respective mode of action has been well-explored in medical sciences (Beyoğlu et al., 2020; Zhang et al., 2020) or in ecotoxicology laboratory-based studies on aquatic models (Le Manach et al., 2018; Huang et al., 2017; Ekman et al., 2015; Gil-Solsona et al., 2017), such methods have been used only faintly in field research so far (Bundy et al., 2009; Lankadurai et al., 2013). Apart from a limited number of evidence of the utility of NMR-based metabolomics in environmental fish studies (Capello et al., 2016; Sotton et al., 2019; Wei et al., 2018), few other examples indicate that field-based LC-MS metabolomics constitutes an emerging and powerful, but still underused, approach for increasing our understanding of *in situ* biological, physiological, ecological or ecotoxicological processes (Meador et al., 2020; Reverter et al., 2017; Goode et al., 2020).

Our analysis constitutes one of the first attempts to push forward the potential of high-throughput molecular phenotyping, and especially through LC-MS-based metabolomics, for environmental assessment. Indeed, this organism molecular phenotyping supported by multi-variable chemometric investigation offers remarkably rich biological information that serves at describing specific phenotypic plasticity. Moreover, this organism variability/responsiveness can further be confronted to local environmental factors such as noxious cyanobacterial metabolites in order to demonstrate correlation or causality relationships.

As described by Pompfret and co-workers (2019), environmental metabolomics exhibit very promising perspectives for operational bio-monitoring applications, according to its reliability, its reproducibility, and its high predictive potential. However, these authors also point out that the responsiveness and robustness of the bio-indicative object, which is characterized through the analytical prism of the metabolomics, remains crucial and have to be carefully evaluated and tested with an appropriate experimental design (Pompfret et al., 2019). However, for ethical concerns, fish bio-monitoring would also gain at been less invasive and deleterious for the organisms. To this end, non-lethal mucus sampling has been investigated by LC-MS and have demonstrated the remarkable informativeness of this approach for environmental studies (Reverter et al. 2017), but these efforts still remain explorative.

## 4. Conclusions

The main challenge facing us remains to be able to develop more integrative approaches that aim at connecting chemical, biological and ecological assessment, in the context of anthropized natural environments confronting multi-stressor pressures. A major caveat of the use of fish environmental metabolomics remains maybe the lack of dedicated databases (Viant et al., 2019) fulfilled by studies considering together different species, populations, development stages, seasons and environments. These data could be used to provide baseline reference values that would further support machine learning or artificial neural network tools for the training of decision-making models.

## Author contributions

B.M. designed experiment, and performed sample analysis. B.M. and A.G. analysed data.

B.M. wrote the manuscript. All authors have given approval to the final version of the manuscript.

## Notes

The authors declare not conflict of interest

## ACKNOWLEDGMENTS

We would like to thanks B. Sotton for his collaboration on NMR data acquisition and treatment. We are also greatful to A. Paris for its support and prolific discussions. This work was supported by grants from CNRS (Défi ENVIROMICS “Toxcyfish” project). The NMR and the MS spectra were respectively acquired at the Plateau technique de Résonance Magnétique Nucleaire and the Plateau technique de spectrométrie de masse bio-organique, Muséum National d’Histoire Naturelle, Paris, France. This work benefitted from the French GDR “Aquatic Ecotoxicology” framework which aims at fostering stimulating scientific discussions and collaborations for more integrative approaches.

## Notes

### Competing Interest Statement

The authors have declared no competing interest.

