## supplementary materials for "Fish metabolome from sub-urban lakes of the Paris area (France) and potential influence of noxious metabolites produced by cyanobacteria"

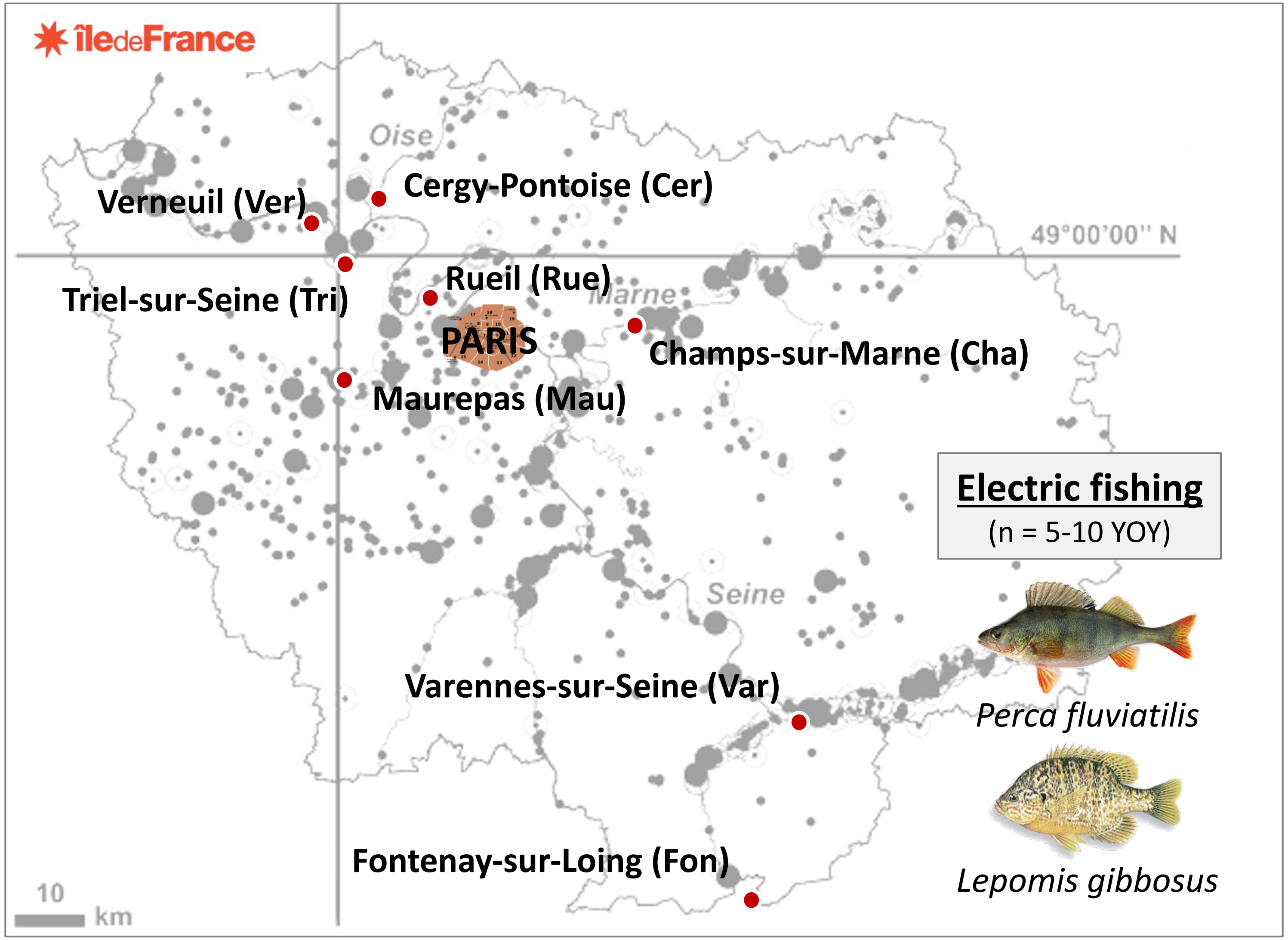


**Supp. fig. S1:** lake map of fish sampling on September 07-10th 2015

| Species/lakes | Cergy-Pontoise | Champs-sur-Marne | Fontenay-sur-Loing | Maurepas | Reuil-Malmaison | Triel-sur-Seine | Varennes-sur-Seine | Verneuil |
|---|---|---|---|---|---|---|---|---|
| *Leucaspius delineatus* | | | 1 | | | | | |
| *Anguilla anguilla* | 11 | 3 | 4 | | | 4 | | 7 |
| *Esox Lucius* | 19 | | | | | | | |
| *Brama sp.* | | | 5 | | | 40 | | 37 |
| *Rhodeus amarus* | | | 73 | 357 | | | | 42 |
| *Cyprinus carpio* | 4 | | | 154 | | 11 | | |
| *Squalius cephalus* | | | 4 | | | | | |
| *Rutilus rutilus* | 1 | | 1 | 1 | 7 | | | 56 |
| *Gobio gobio* | | | | 5 | | | | |
| *Gymnocephalus cernuus* | | 5 | | 15 | | | | 3 |
| ***Perca fluviatilis*** | **10** | **26** | **37** | **27** | **26** | **20** | **9** | **25** |
| ***Lepomis gibbosus*** | **368** | **358** | **19** | **57** | **6** | **17** | | **42** |
| *Ameiurus melas* | 1 | | | | | | | |
| *Scardinius erythrophthalmus* | 21 | | 23 | 12 | 16 | 52 | | 2 |
| *Silurus glanis* | | | 2 | | | | | 5 |
| *Tinca tinca* | 15 | 42 | | 3 | | 6 | | |
| Total number of species | 10 | 5 | 10 | 8 | 4 | 7 | 1 | 9 |
| Total number of specimens | 450 | 434 | 169 | 631 | 55 | 150 | 9 | 219 |

**Supp. table S1.** Effectives of fish collection in the 8 lakes sampled on September 7-10th 2015.

| Pond | Localisation | Year of study | Surface area (ha) | Max depth (m) | Mean dissolved O2 concentration (% sat.) | Mean pH | Mean conductivity (µS.cm$^{-1}$) | Mean Temperatures (°C) | Total Chl *a* concentration (µg.L$^{-1}$ eq. Chl *a*) | Total MC concentration (µg.L-1 eq. MC-LR) |
|---|---|---|---|---|---|---|---|---|---|---|
| **Cer** | 49°01’ N 02°03’ E | 2015 | 10.6 | 5 | 78.5 ± 18.7 | 8.3 ± 0.3 | 270.1 ± 6.9 | 18.8 ± 0.2 | 26.7 ± 1.3 | 0.08 ± 0.1 |
| **Cha** | 48°51’ N 02°35’ E | 2015 | 10.3 | 3.3 | 42 ± 11.4 | 7.8 ± 0.2 | 497.9 ± 5.2 | 18.5 ± 0.3 | 52.6 ± 1 | 0 ± 0.1 |
| **Fon** | 48°06’ N 02°75’ E | 2015 | 22 | 6 | 59.4 ± 5.4 | 7.5 ± 0.2 | 387.7 ± 2.3 | 19.5 ± 0.1 | 112.7 ± 6 | 0.03 ± 0.1 |
| **Mau** | 48°46’ N 01°55’ E | 2015 | 7.8 | 5.8 | 65 ± 42 | 7.1 ± 0.6 | 273.7 ± 102.2 | 16.4 ± 0.9 | 102.4 ± 2.8 | 0.04 ± 0.1 |
| **Rue** | 48°51’ N 02°10’ E | 2015 | 1.4 | 3.7 | 40.3 ± 1.4 | 7.6 ± 0.1 | 604.8 ± 10.7 | 16.6 ± 0.2 | 82.4 ± 3.7 | 0 ± 0.1 |
| **Tri** | 48°57’ N 02°00’ E | 2015 | 10 | 3 | 81.7 ± 31.9 | 9.2 ± 0.4 | 862.2 ± 13.6 | 18.2 ± 0.4 | 41.1 ± 1.9 | 2.07 ± 0.1 |
| **Ver** | 49°00’ N 01°58’ E | 2015 | 46.2 | 4.6 | 84.2 ± 15.2 | 9.1 ± 0.1 | 288.6 ± 19.5 | 19.1 ± 0.1 | 137 ± 5.4 | 0.17 ± 0.1 |
| **Var** | 48°22’ N 02°57’ E | 2015 | 16.5 | 4.8 | 102.1 ± 16.5 | 9.2 ± 0.1 | 293.8 ± 2.1 | 18.3 ± 0.4 | 75.1 ± 4.3 | 3.37 ± 0.1 |

**Supp. table S2.** Descriptors and metadata of the 8 lakes

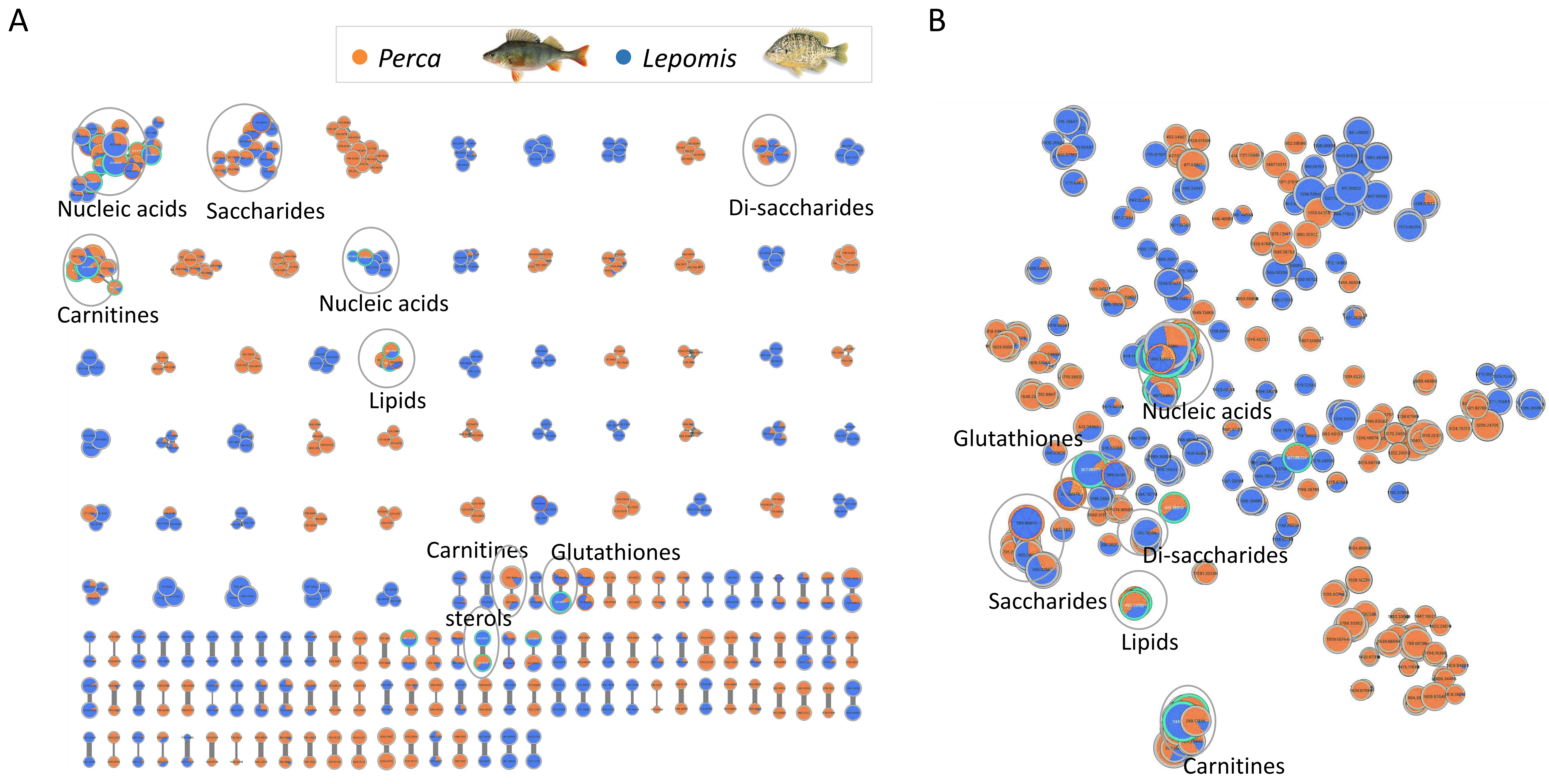


**Supp. fig. S2:** Molecular networking of *Perca* and *Lepomis* LC-MS/MS metabolomes characterized by GNPS (A) and *t*-SNE (B) algorithms.

**Supp. fig. S3:** Score plot (A) and loading plot (B) of a PCA and heatmap with hierarchical clustering performed with LC-MS data from the phytoplanctonic biomass collected from the water column of 8 sampled lakes.

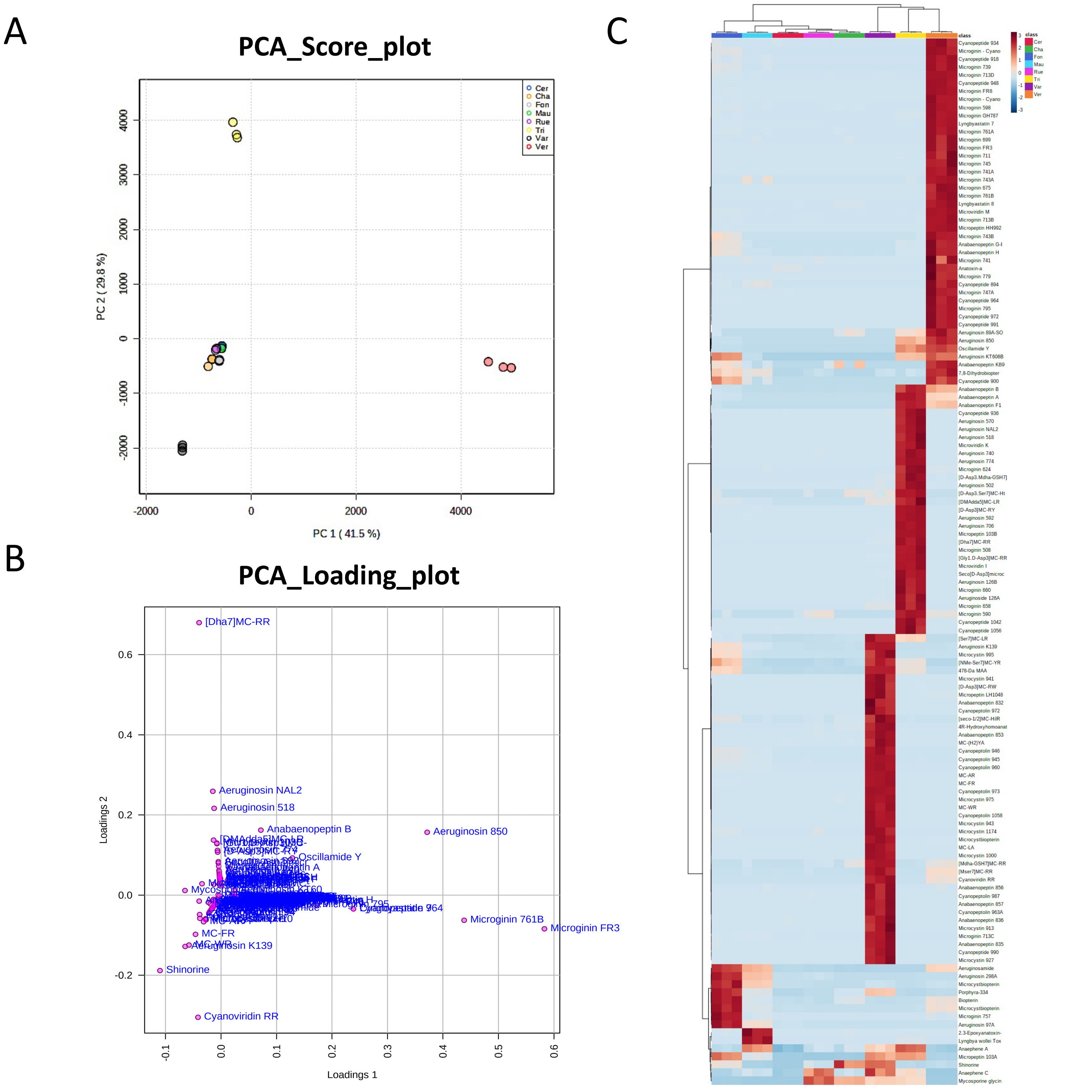

A B

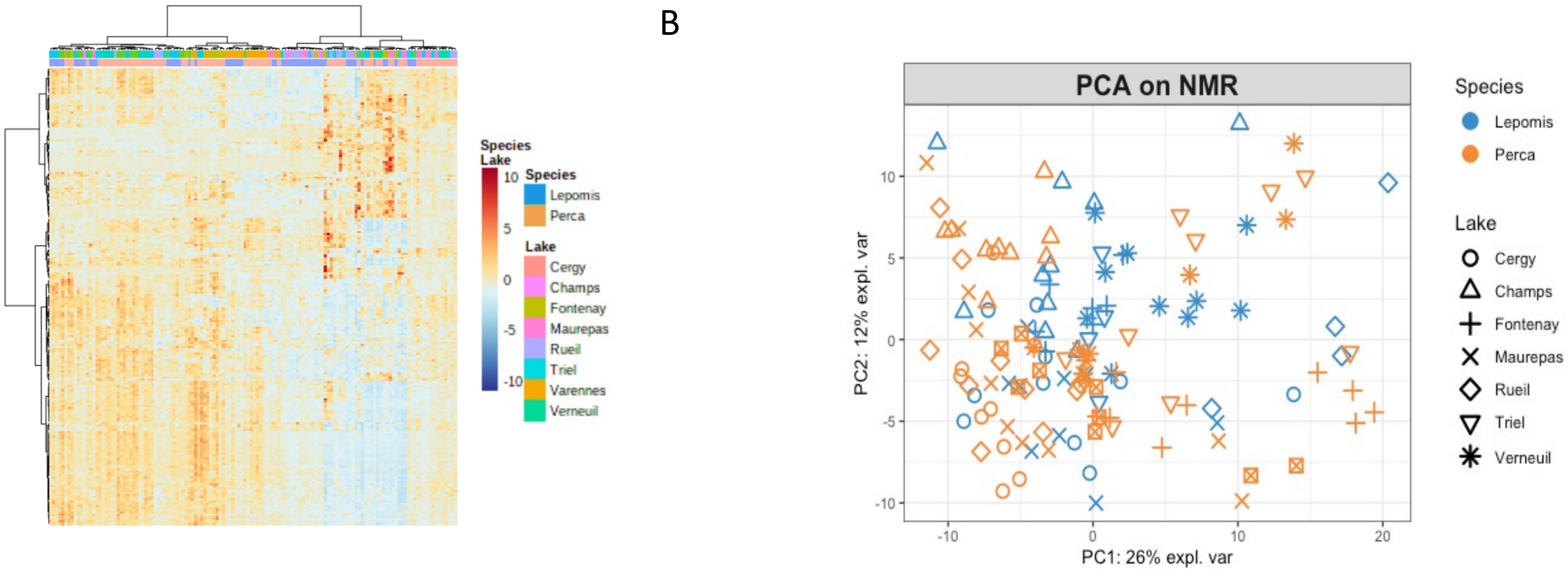


**Supp. fig. S4:** $^{1}$H NMR metabolomics of *Perca* and *Lepomis* livers illustred by heatmap with hierarchical classification (A) and PCA (B).

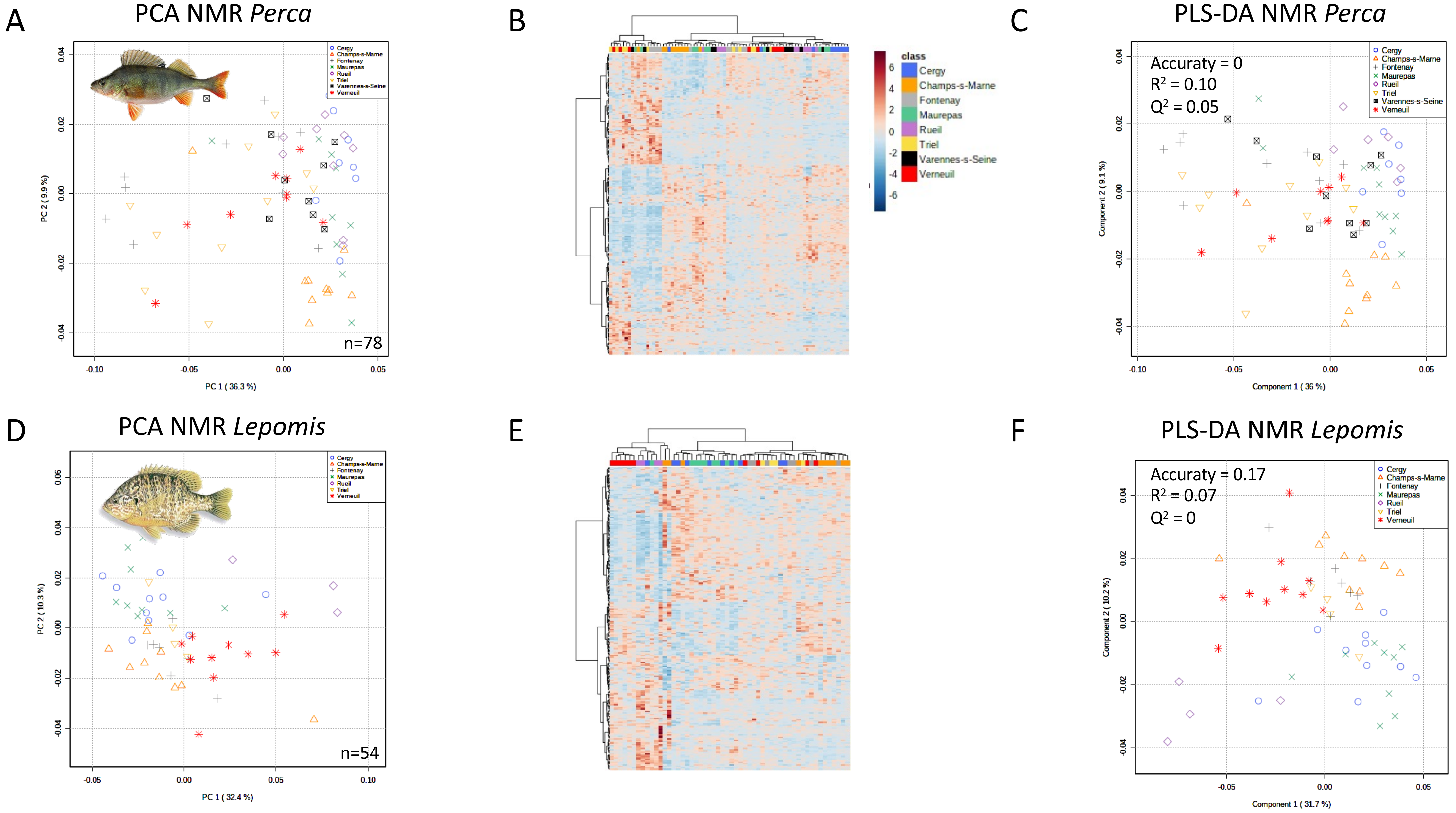


**Supp. fig. S5:** $^{1}$H NMR metabolomics of *Perca* (A-C) and *Lepomis* (D-F) livers illustred by PCA (A and D), heatmap with hierarchical classification (B and E) and PLS-DA regarding the lake of sampling (C and F).

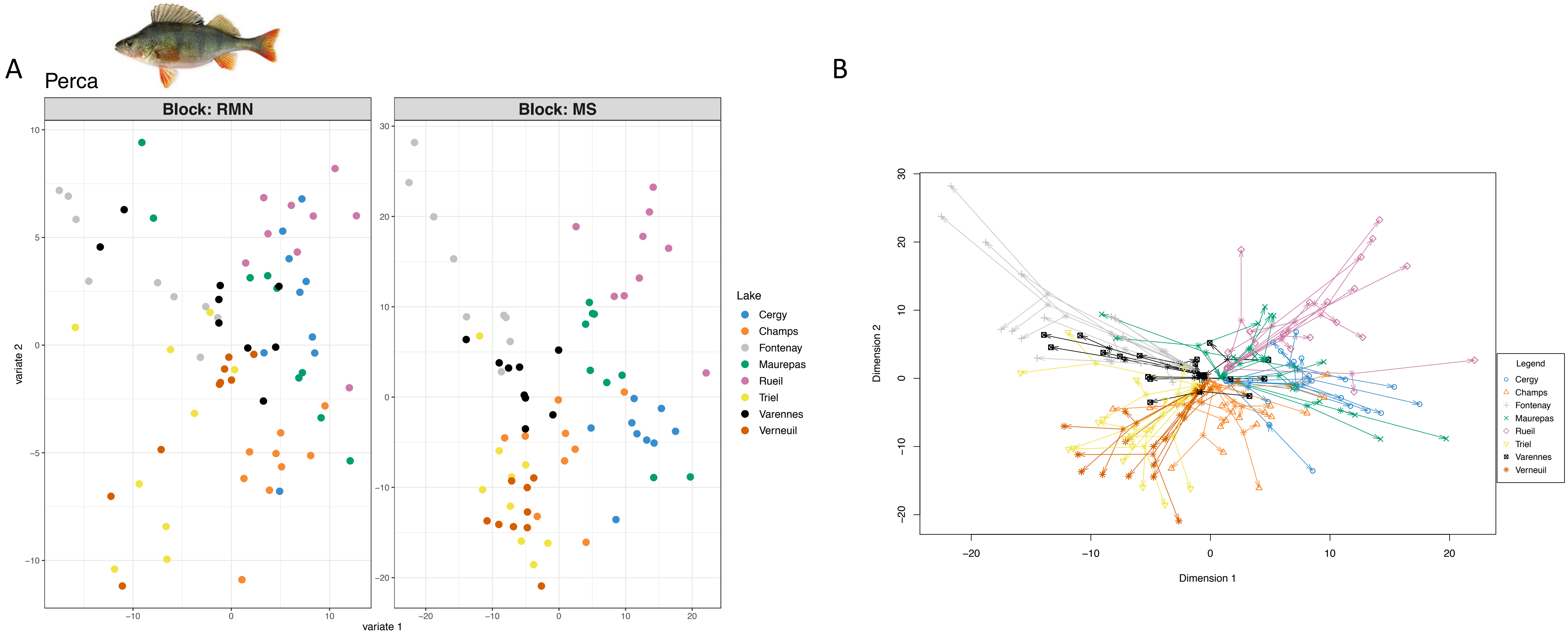


**Supp. fig. S6:** Distinct (A) and merged (B) PLS-DA of the integrated NMR and LC-MS datasets of *Perca* liver metabolome using MixOmics (global correlation score = 0.79).

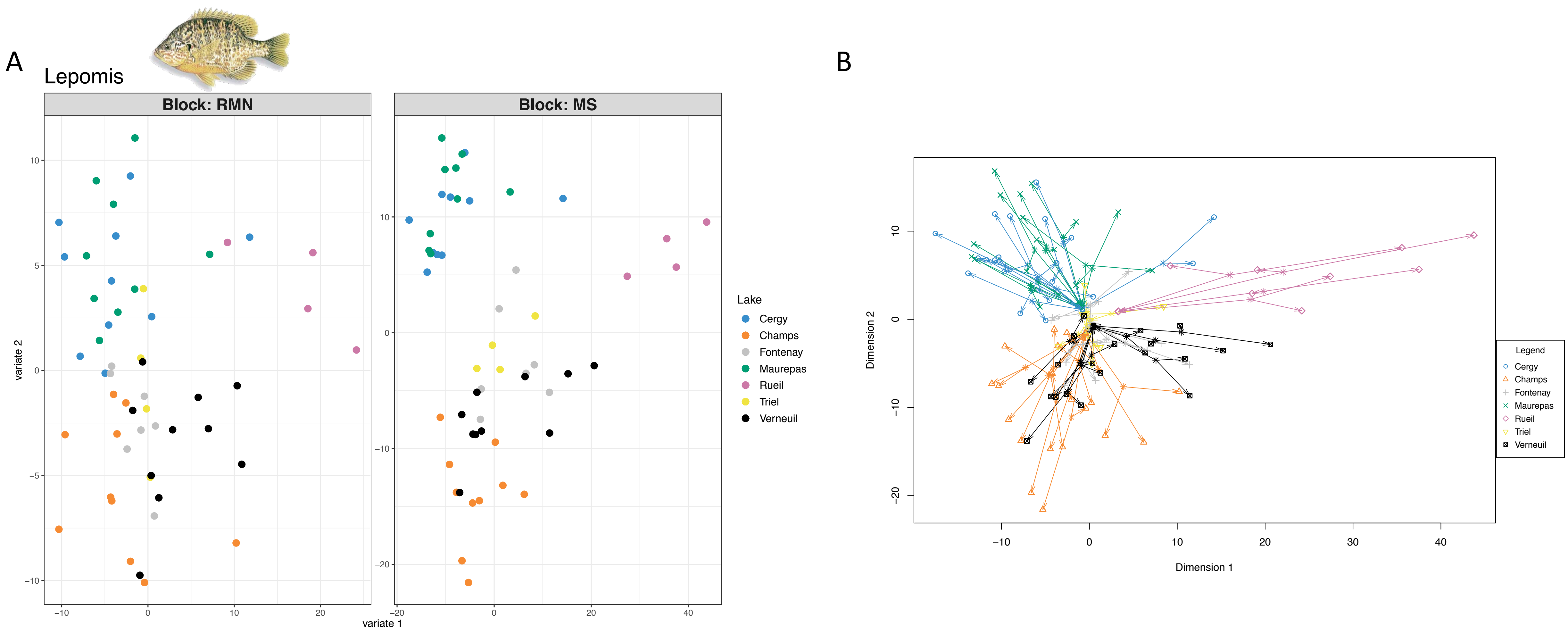


**Supp. fig. S7:** Distinct (A) and merged (B) PLS-DA of the integrated NMR and LC-MS datasets of *Lepomis* liver metabolome using MixOmics (global correlation score = 0.83).
